# MetaSage: Machine Learning-Based Prioritization of Metabolic Regulators from Multi-Omics Data

**DOI:** 10.1101/2025.07.09.663994

**Authors:** Chenwei Wang, John M. Elizarraras, Bing Zhang

## Abstract

Dysregulation of metabolites is a hallmark of cancer, yet the underlying regulatory mechanisms remain poorly understood. To systematically explore metabolic regulation across cancers, we developed an XGBoost-based machine learning pipeline, MetaSage, that integrates context-agnostic knowledge graph with multi-omics datasets. Using harmonized data from 15 cohorts spanning 11 cancer types, we identified 442 variable metabolites and found that both genes and upstream metabolites showed comparable regulatory influence. Predictable metabolites, defined by a significant correlation between predicted and measured levels, were identified using our pipeline and varied widely across cohorts-partially due to the batch effect. For each predictable metabolite, key regulatory features were determined using Shapley values. This yielded 1,146 gene features and 363 precursor metabolites as important regulators. Network analysis of 22 recurrent metabolites revealed a mix of conserved and cancer type-specific regulatory patterns. Our framework enables robust discovery of metabolite regulation and therapeutic insights in cancer.

## Introduction

Metabolites, the small-molecule intermediates and end products of cellular metabolism, play crucial roles in cancer biology^1–3^. In addition to fueling biomass synthesis and energy production, they participate in the regulation of gene expression, signal transduction, and epigenetic modifications that drive tumor initiation and progression^1^. Aberrant metabolic reprogramming is now recognized as a hallmark of cancer, enabling malignant cells to adapt to fluctuating nutrient conditions and sustain rapid proliferation^3^. One prominent example is the Warburg effect—also known as aerobic glycolysis—which represents a characteristic metabolic shift in many cancer cells^4^. This adaptation allows cancer cells to metabolize glucose at high rates and produce lactate even in the presence of oxygen, thereby supporting the biosynthetic and energetic demands necessary for rapid growth, survival, and invasion. Such metabolic rewiring is driven not only by nutrient availability—which influences the levels of upstream metabolites feeding into metabolic pathways—but also by the activity of associated enzymes, transporters, and signaling proteins, many of which are dysregulated in cancer. For example, hexokinase 2 (HK2) and pyruvate dehydrogenase kinase 1 (PDK1) are key regulators that promote the Warburg effect^4^. This intricate interplay between metabolism and its protein regulators underscores the therapeutic potential of targeting metabolic vulnerabilities in oncology and highlights the need for a deeper understanding of metabolic regulation in tumor development and progression^5^.

Over the past decades, advances in high-throughput analytical technologies, such as mass spectrometry (MS), together with the refinement of metabolite libraries, have propelled the development of metabolomics and enabled the systematic analysis of metabolic regulation and the identification of therapeutic opportunities across various cancer types^6–9^. Increasingly, studies are integrating metabolomics with other omics platforms such as transcriptomics and proteomics, thereby providing valuable resources to investigate the co-regulation of metabolites and their associated genes^9,10^. However, most multi-omics analyses to date have focused on indirect associations or statistical correlations between metabolite levels and gene expression. Methods capable of systematically inferring direct regulatory interactions between genes and metabolites remain limited, representing a major challenge and opportunity in the field.

In this study, we developed a machine learning–based pipeline named MetaSage to systematically investigate the regulation of cancer-associated metabolites. By integrating established context agnostic metabolite–gene knowledge graphs with disease-specific multi-omics datasets that include metabolomic profiles from various cancer cohorts, our approach enables the identification of both conserved and cancer type–specific regulatory mechanisms. For instance, we identified consistent regulatory patterns for metabolites such as adenine and guanine across multiple cancer types, while also revealing cancer type-specific regulatory influences on the metabolites involved in energy metabolism in distinct tumor contexts.

Taken together, our method offers a novel and systematic framework to decipher metabolic regulation in diverse cancer settings and serves as a valuable tool to support mechanistic studies and biomarker discovery in oncology.

## Results

### Overview of MetaSage pipeline

MetaSage is a machine learning pipeline designed to identify study-specific key regulators of variable metabolites by integrating metabolomic profiles with at least one gene-level dataset (e.g., RNA-seq or proteomics) from the same cohort. Built on the XGBoost algorithm, MetaSage comprises three main steps:

### Feature Selection (Figure 1A)

Variation in metabolite abundance is influenced by the availability of upstream precursors and the activity of enzymes involved in its synthesis and degradation. To capture these determining features for each target metabolite, MetaSage extracts all directly associated metabolic reactions from a comprehensive human genome-scale metabolic model (GEM)^11^. From these reactions, it identifies upstream (precursor) metabolites and the corresponding anabolic and catabolic enzymes. This information is then integrated with study-specific multi-omics datasets, including metabolomic profiles and at least one gene-level dataset (e.g., RNA-seq or proteomics). Only features (genes and metabolites) quantified in these datasets are retained for downstream analysis.

**Figure 1.**
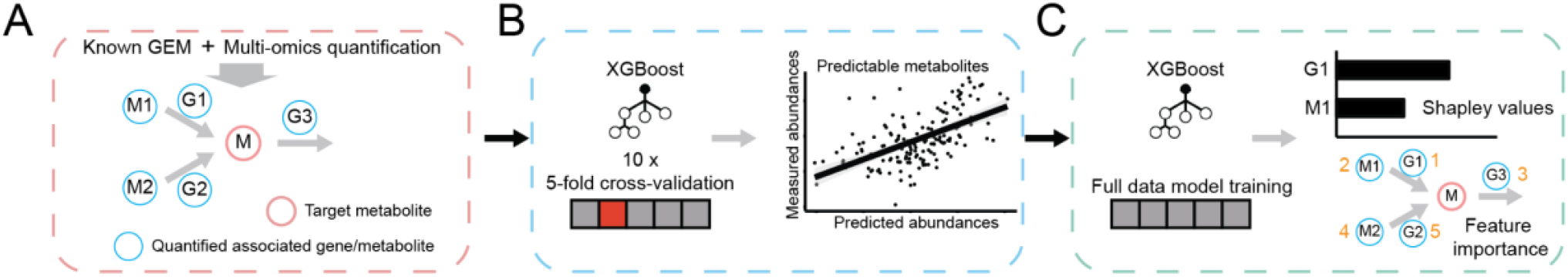
Overview of the MetaSage pipeline. The MetaSage pipeline consists of 3 key steps: (A) Feature Selection: For each target metabolite, associated genes and precursor metabolites were selected based on human GEM and multi-omics quantification. (B) Predictability Assessment: XGBoost regression models were trained using 10 iterations of 5-fold cross-validation. Metabolites with a significant Pearson correlation between predicted and measured intensities were defined as predictable. (C) Regulator Prioritization: For each predictable metabolite, models were retrained on the full dataset and feature importance was calculated using Shapley values (SHAP), identifying the top-ranked features as key regulators.

### Predictability Assessment (Figure 1B)

For each target metabolite with at least one quantified feature, MetaSage applies XGBoost to predict its abundance using 10 iterations of 5-fold cross-validation. Pearson correlation coefficients are then calculated between the average predicted metabolite levels and the observed ones. Metabolites showing statistically significant positive correlations exceeding a user-defined correlation threshold are considered to exhibit a predictive signal and classified as informatively predictable.

### Regulator Prioritization (Figure 1C)

For each informatively predictable metabolite, MetaSage retrains the model using the complete dataset. Feature importance is evaluated using Shapley values (SHAP), and features are ranked according to their mean absolute SHAP values. The top-ranking features are considered the most influential regulators of the corresponding metabolite.

### Pan-Cancer Identification of Informatively Predictable Metabolites

Paired metabolomic and gene expression datasets were obtained from a previous pan-cancer study^10^, encompassing harmonized data from 15 cohorts across 11 cancer types. We focused exclusively on tumor samples due to the limited availability of normal samples in most cohorts (**Figure 2A**). As expected, coverage of protein-coding genes was relatively consistent across the RNA-seq datasets (17,494 to 18,919), whereas metabolite quantification varied substantially among cohorts, ranging from 178 to 1,006 quantified metabolites (**Figure 2B**). Reflecting this variability, we observed limited overlap across metabolomics datasets: over 50% of metabolites were uniquely quantified in a single cohort, while more than 75% of genes were consistently covered across all 15 RNA-seq cohorts (**Figure 2C**).

**Figure 2.**
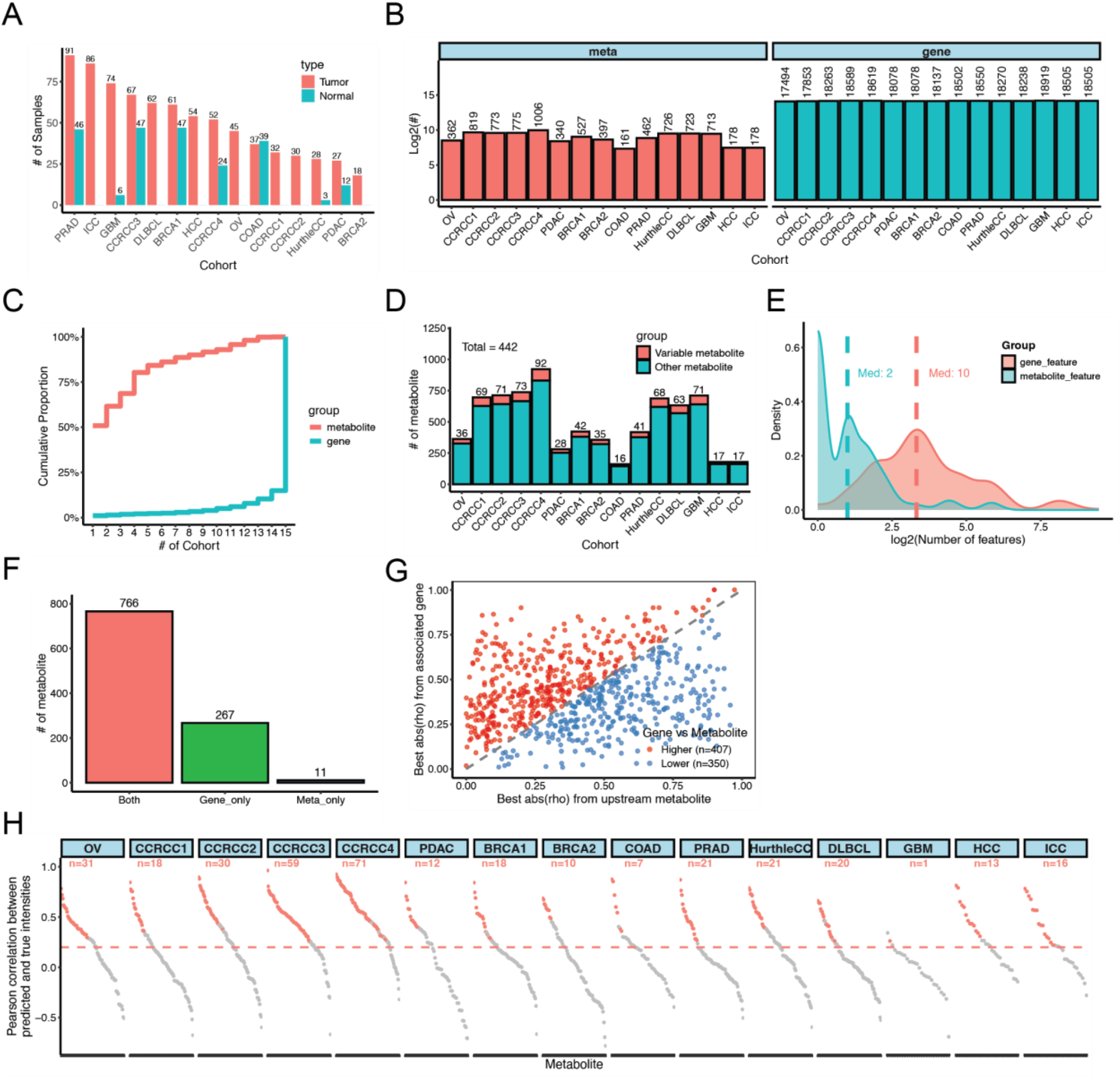
Pan-cancer application of the MetaSage pipeline. (A) Overview of 15 cancer cohorts from 11 cancer types used in this study. (B) Number of quantified metabolites and genes per cohort. (C) Cumulative distribution function curve showing the overlap of quantified metabolites and genes across all cohorts. (D) Top 10 variable metabolites were chosen from each cohort. (E) Distribution of selected gene and metabolite features per target metabolite. Dashed line indicate median counts. (F) Proportion of target metabolites with gene-only, metabolite-only, or both feature types. (G) Comparison of the best Spearman correlation coefficients between target metabolites and their associated gene or precursor features. (H) Number of predictable metabolites (R > 0.2, p < 0.05) identified in each cohort. Red dots represent the predictable metabolites.

To focus on cancer-relevant metabolic alterations while maintaining broad coverage, we selected the top 10% most variable metabolites from each cohort, yielding a set of 442 unique metabolites as input for MetaSage pipeline (**Figure 2D**). The pipeline was then applied independently to each cohort using these metabolites as prediction targets. Through feature selection, we identified substantially more gene features than metabolite features associated with each target metabolite, with median counts of 10 and 2, respectively (**Figure 2E**). Most target metabolites were associated with both gene and metabolite features, and only 11 metabolites were exclusively linked to metabolite features (**Figure 2F**).

To systematically assess the regulatory contribution of associated genes and precursor metabolites, we computed Spearman correlation coefficients between each target metabolite and its associated genes or precursors. For each metabolite, we identified the stronger of the two correlations (absolute rho values) and observed slightly better predictive performance from gene features: 407 metabolites showed stronger correlations with gene features, while 350 showed stronger correlations with precursor metabolites (**Figure 2G**). These results suggest that both gene expression and upstream metabolite levels contribute comparably to metabolite abundance, supporting the integration of both feature types in predictive modeling.

Predictable metabolites were defined in each cohort, with a Pearson correlation threshold of R > 0.2 and p-value < 0.05 between predicted and measured abundances. The number of predictable metabolites varied widely across cohorts, ranging from only 1 in GBM to 71 in CCRCC3. Interestingly, even among cohorts representing the same cancer type (e.g., BRCA or CCRCC), notable differences were observed in the number of predictable metabolites identified (**Figure 2H**), suggesting that sample size, data quality, or tumor heterogeneity may influence predictability.

### Feature Importance Evaluation of Predictable Metabolites

For each predictable metabolite, feature importance was assessed by retraining the model using all samples within the cohort. The top contributing features were then identified based on their average absolute Shapley values. Specifically, in addition to the top-ranked feature, any other features with at least 10% of the top feature’s contribution were retained. A maximum of five features were selected per metabolite. Across all 15 cohorts, this process yielded a total of 1,146 important gene features (75.9%) and 363 precursor metabolites (24.1%) (**Figure 3A**).

**Figure 3.**
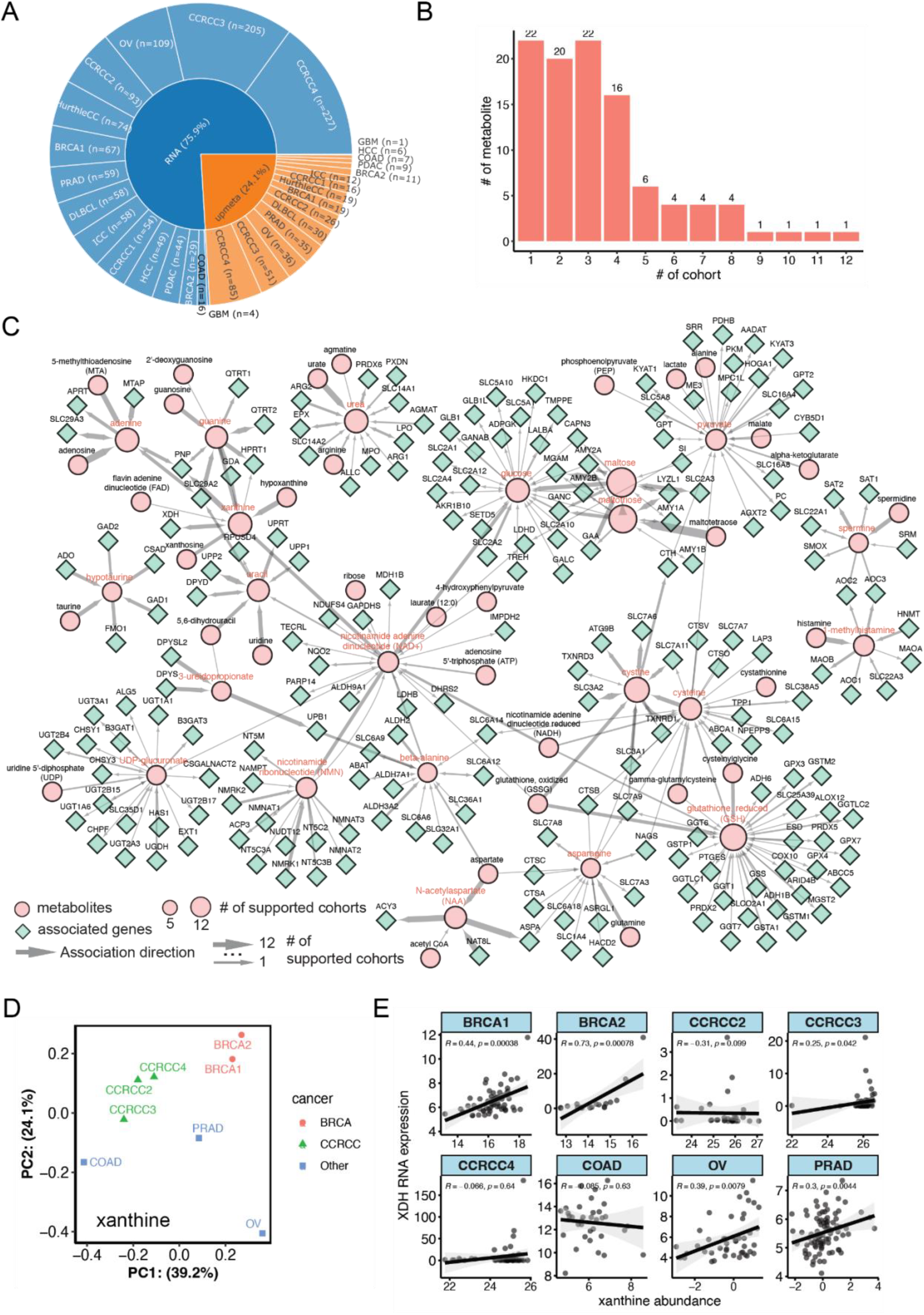
Prioritization and network analysis of key regulatory features for predictable metabolites. (A) Distribution of the important features across 15 cohorts. (B) Number of cohorts in which each predictable metabolite was identified. (C) Network connecting the 22 recurrent metabolites to their important features. Target metabolites are colored in red, with node size indicating the number of cohorts in which they were identified. Edge width reflects the number of cohorts where a given feature was identified as important, arrows indicate their relationship with target metabolites (upstream, downstream, or bidirectional). (D) PCA plot showing the distribution of 8 cohorts in which xanthine was identified as a predictable metabolite, colored by cancer type: BRCA (red), CCRCC (green) and Other cancer types (blue). (E) Spearman’s correlations between xanthine abundance and RNA expression of XDH in 8 cohorts with xanthine identified as predictable metabolites

Among all predictable metabolites, 22 were identified in at least five cohorts (**Figure 3B**), offering a valuable opportunity to explore their regulatory mechanisms across different cancer types. To visualize this, we constructed a regulatory network including these 22 metabolites and their corresponding important features (**Figure 3C**). In this network, edge width reflects the number of cohorts in which a given feature was identified as important. We found that precursor metabolites were consistently identified as key features across cohorts, as indicated by the generally thicker edges.

Interestingly, the important features for some metabolites—such as glucose and Nicotinamide Adenine Dinucleotide (NAD+)—varied widely across cancer types, suggesting divergent regulatory mechanisms. In contrast, other metabolites, such as N-acetylaspartate (NAA) and adenine, exhibited more conserved regulatory patterns with similar key features across multiple cancer types. This consistency may be partially attributed to the limited number of mapped features (**Figure 3C**).

To further evaluate the robustness and specificity of MetaSage, we compared prediction results across different cancer types. For predictable metabolites shared by multiple cohorts, we performed principal component analysis (PCA) based on the average absolute Shapley values of their input features. Cohorts from the same cancer type tended to cluster together. For example, xanthine was identified as a predictable metabolite in eight cohorts, with the three CCRCC cohorts clustering together and the two BRCA cohorts forming a separate cluster (**Figure 3D**). Similar clustering patterns were observed for glucose and maltose (**Figure S1A, B**). These results suggest that cohorts from the same cancer type tend to share similar key predictive features, while these features may differ across cancer types. For example, in BRCA, XDH—an enzyme that converts hypoxanthine to xanthine—was identified as the top-ranked predictive feature for xanthine abundance, whereas it was not among the top features in CCRCC. Instead, hypoxanthine was predicted as a top feature in one CCRCC cohort and important feature in other two CCRCC cohorts. Supporting these predictions, XDH showed strong positive RNA–metabolite correlation with xanthine in both BRCA cohorts, but weak correlations in the CCRCC cohorts (**Figure 3E**). In contrast, hypoxanthine showed strong positive correlations with xanthine in the CCRCC cohorts but not the BRCA cohorts (**Figure S1C**). These findings suggest that xanthine regulation in BRCA may be primarily driven by XDH expression, while in CCRCC it may depend more on hypoxanthine availability or upstream pathway activity.

Together, these results demonstrate that our machine learning pipeline, when integrated with multi-omics data, can effectively identify key regulators of metabolite abundance and provide insight into both conserved and cancer type-specific regulatory mechanisms.

## Discussion

Understanding metabolic regulation across different cancer types remains a significant challenge. Over the past decades, numerous efforts have been devoted to uncovering the underlying mechanisms from various perspectives. For instance, genome-scale metabolic models (GEMs)^11^ have been developed to systematically map known metabolic reactions and their associated genes, providing a comprehensive framework for exploring metabolic networks. In parallel, experimental approaches such as ^13^C metabolic flux analysis offer precise, pathway-specific insights by tracing labeled substrates through metabolic pathways^12^. Metabolomics has also emerged as a powerful tool for profiling thousands of metabolites simultaneously, offering a biochemical snapshot of cellular states^6^. However, despite these advancements, metabolite concentrations often serve only as indirect proxies for pathway activity due to the highly dynamic and context-dependent nature of metabolic fluxes^12^. This limitation underscores the need for integrative strategies that combine metabolomics with other omics layers—such as transcriptomics and proteomics—to provide a more holistic, systems-level view of metabolic regulation.

Traditionally, metabolic pathway analysis relies on identifying differentially abundant metabolites and genes between tumor and normal tissues, followed by enrichment analysis to infer affected pathways^13^. While informative for detecting general metabolic shifts, such methods fall short in revealing the key regulatory mechanisms that drive these changes. To address this gap, we present MetaSage, a computational framework that integrates metabolomics with multi-omics data to identify regulators underlying cancer-associated metabolic rewiring. By leveraging machine learning, MetaSage identifies the most influential regulators based on their contribution to predictive models, providing an interpretable and data-driven approach to uncover metabolic control points. When applied to diverse cancer cohorts, MetaSage revealed substantial inter-cohort variability. Nevertheless, we achieved reliable coverage and consistent identification of regulatory features for several critical metabolites, such as xanthine and glucose, which exhibited cancer-type-specific regulatory patterns. Conversely, other metabolites showed more conserved regulation across cancers, suggesting shared metabolic mechanisms. These findings provide valuable insights into both common and context-specific regulatory architectures and offer promising leads for therapeutic targeting.

There are, however, several avenues for further improvement. One limitation is the current reliance on transcriptomic data as the primary gene-centric omics layer. Incorporating additional data types, such as proteomics and post-translational modification (PTM) profiles, could enhance the accuracy and depth of regulatory inference. This remains constrained by data availability, as few cohorts currently offer comprehensive multi-omics coverage. Additionally, even with detailed GEMs, certain reactions may be underrepresented if their associated components are missing from the input data, potentially causing regulators from secondary pathways to appear disproportionately important. This limitation could be mitigated by integrating flux-based approaches^14^, which leverage gene expression and provide pathway-level activity estimates, allowing prioritization of biologically relevant pathways. Moreover, MetaSage currently focuses on direct gene–metabolite relationships. This design may overlook longer-range regulatory influences, especially when intermediate genes or metabolites are not captured in the dataset. Future versions of the framework should incorporate more extensive pathway topology from GEMs to infer indirect regulatory links, while also carefully managing the added complexity during interpretation.

In conclusion, MetaSage offers a novel and interpretable approach to dissect metabolic regulation by integrating metabolomics with multi-omics data. While further improvements are needed to enhance resolution and generalizability, our study demonstrates the power of combining computational modeling and machine learning to decode the regulatory logic of cancer metabolism.

## Methods

### Pan-cancer metabolomic and gene expression data

Reprocessed and normalized metabolomic and gene expression data from 15 cancer cohorts were downloaded from https://zenodo.org/records/7874228, along with the ID mapping table for quantified metabolites. For each cohort, only samples with both metabolomic and gene expression data were retained for downstream analysis. Metabolites and genes with all zero values were excluded.

### MetaSage pipeline

The human genome-scale metabolic model (GEM) was obtained from the Metabolic Atlas (https://metabolicatlas.org/, version 1.14.0)11, which includes 13,024 metabolic reactions involving 8,363 metabolites and 2,920 genes. Metabolites identified in this study were mapped to the GEM using the human metabolome database (HMDB)15 IDs.

For each target metabolite quantified in at least 10 samples, we extracted all associated genes from the metabolic reactions in which the metabolite is involved. Only genes with available expression data in the transcriptomic dataset were retained. Gene expression levels of these genes were used as input features. Additionally, upstream (precursor) metabolites detected in the metabolomics dataset were included as extra features.

For each target metabolite with at least one input feature, an XGBoost regression model was trained to predict metabolite intensity based on the selected features. Five-fold cross-validation was repeated 10 times to ensure robust prediction. Pearson correlation coefficients were calculated between the average predicted and measured metabolite intensities. Metabolites with a correlation coefficient (R) > 0.2 and a p-value < 0.05 were defined as predictable.

For each predictable metabolite, a final model was trained on the full dataset still using XGBoost, with Shapley values computed during training to assess feature importance. The average absolute Shapley value was calculated for each feature. Default XGBoost parameters were used, except for n_estimators = 100 and random_state = 1 through 10 for the repeated cross-validations.

For network visualization, associated genes were categorized into three groups based on the reactions they participated in:

1. **Upstream genes**: genes involved in reactions that produce the target metabolite;
2. **Downstream genes**: genes involved in reactions that consume the target metabolite;
3. **Bi-directional genes**: genes involved in reversible reactions involving the target metabolite.

In addition to the top-ranked feature with the highest average absolute Shapley value, other features were retained if their contribution was at least 10% of that of the top feature. For each metabolite, a maximum of five features were selected.

## Acknowledgements

This study was supported by the National Cancer Institute (NCI) CPTAC award U24 CA271076 (B.Z.) and funding from the McNair Medical Institute at The Robert and Janice McNair Foundation (B.Z.).

## Authors Contributions Statement

B.Z., and C.W. conceived the study. C.W. developed the pipeline and performed data analysis. J.M.E built the website. C.W., and B.Z. interpreted the data and wrote the manuscript. All authors reviewed and approved the manuscript.

## Code Availability

The source code of MetaSage is available upon request.

## Figure legends

**Figure S1.**
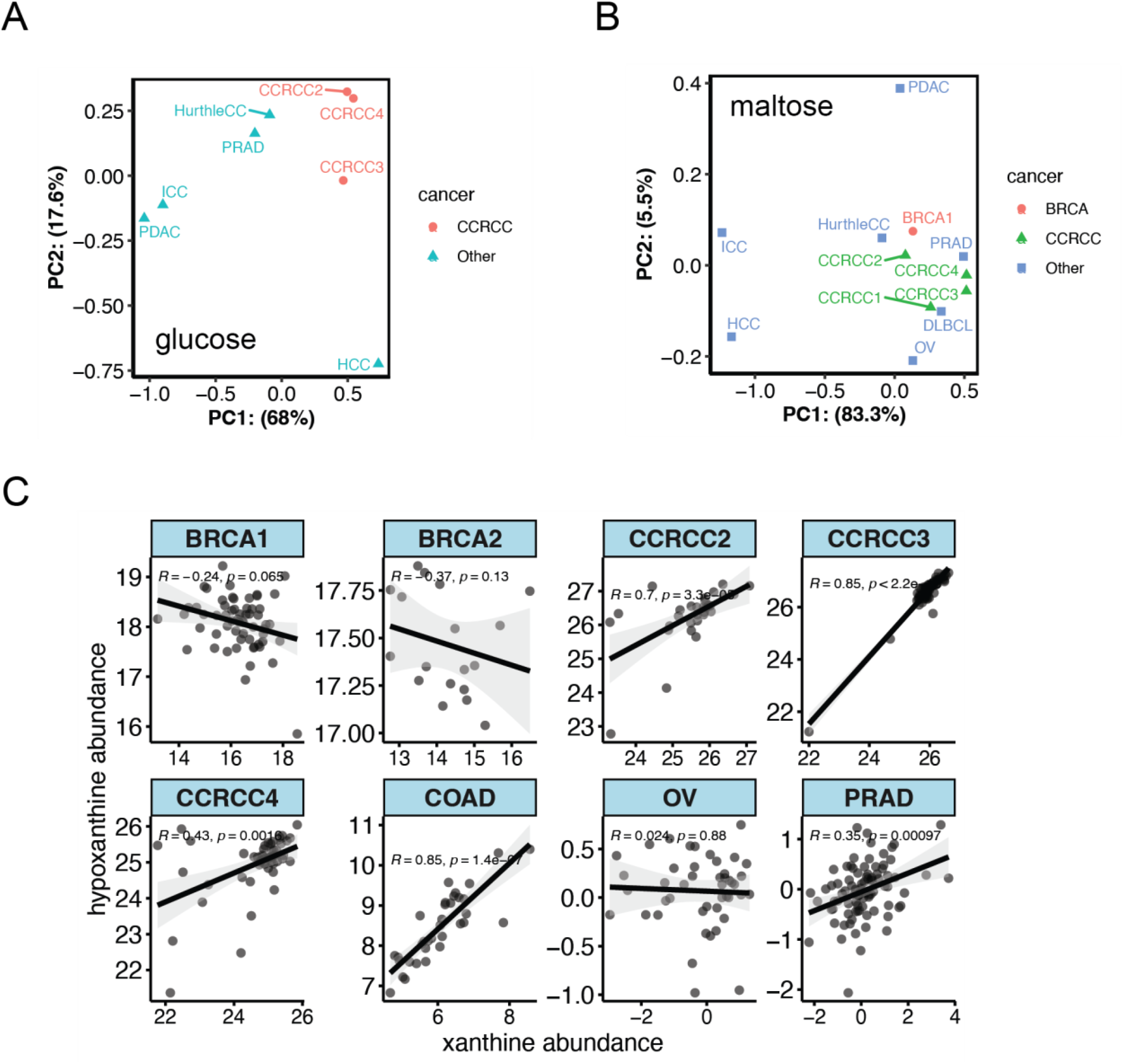
Additional analysis of predictable metabolites shared across multiple cohorts. (A) PCA plot showing the distribution of 8 cohorts in which glucose was identified as a predictable metabolite, colored by cancer type: CCRCC (red) and Other cancer types (blue). (B) PCA plot showing the distribution of 12 cohorts in which maltose was identified as a predictable metabolite, colored by cancer type: BRCA (red), CCRCC (green) and Other cancer types (blue). (C) Spearman’s correlation between xanthine and hypoxanthine abundance in 8 cohorts where xanthine was identified as a predictable metabolite.

